# Multi-omics profiling of CHO parental hosts reveals cell line-specific variations in bioprocessing traits

**DOI:** 10.1101/532150

**Authors:** Meiyappan Lakshmanan, Yee Jiun Kok, Alison P. Lee, Sarantos Kyriakopoulos, Hsueh Lee Lim, Gavin Teo, Swan Li Poh, Wen Qin Tang, Jongkwang Hong, Andy Hee-Meng Tan, Xuezhi Bi, Ying Swan Ho, Peiqing Zhang, Say Kong Ng, Dong-Yup Lee

**Author notes:** Correspondence to: D.-Y. Lee, Contract grant sponsor: Agency for Science, Technology and Research, Singapore, Contract grant sponsor: Next-Generation BioGreen 21 Program, Rural Development Administration, Republic of Korea, Contract grant number: PJ01334605.

## Abstract

Chinese hamster ovary (CHO) cells are the most prevalent mammalian cell factories for producing recombinant therapeutic proteins due to their ability to synthesize human-like post-translational modifications and ease of maintenance in suspension cultures. Currently, a wide variety of CHO host cell lines have been developed; substantial differences exist in their phenotypes even when transfected with the same target vector. However, relatively less is known about the influence of their inherited genetic heterogeneity on phenotypic traits and production potential from the bioprocessing point of view. Herein, we present a global transcriptome and proteome profiling of three commonly used parental cell lines (CHO-K1, CHO-DXB11 and CHO-DG44) in suspension cultures and further report their growth-related characteristics, and N- and O-glycosylation patterns of host cell proteins (HCPs). The comparative multi-omics analysis indicated that some physiological variations of CHO cells grown in the same media are possibly originated from the genetic deficits, particularly in the cell cycle progression. Moreover, the dihydrofolate reductase deficient DG44 and DXB11 possess relatively less active metabolism when compared to K1 cells. The protein processing abilities and the N- and O-glycosylation profiles also differ significantly across the host cell lines, suggesting the need to select host cells in a rational manner for the cell line development on the basis of recombinant protein being produced.

## 1 INTRODUCTION

Mammalian expression systems are the preferred choice for the industrial production of recombinant glycoprotein therapeutics due to their humanized post-translational modifications, i.e. N-glycosylation and appropriate protein folding, and the ability to secrete the product naturally outside of the cell (Wurm, 2004). Among several mammalian host cells, Chinese hamster ovary (CHO) cells are commonly used since they can grow in suspension culture with chemically defined media, and are highly resistant to viral susceptibility. Overall, 84% of the biopharmaceutical drugs are currently produced using CHO cells (Walsh, 2018).

Generally, bioprocess development includes two major stages: 1) establishment of efficient cell lines producing target proteins, and 2) development of optimal cell culture media and/or process conditions (Hong et al., 2018). Currently, a multitude of CHO cells with various genetic backgrounds have been utilized since the first approval of a drug produced in CHO cells in 1984 (Kaufman et al., 1985). Of them, the CHO-K1, -DXB11 and -DG44 are the most relevant parental cell lines to manufacture target products industrially (Golabgir et al., 2016). Historically, the proline-deficient K1 cells was established in 1968 (Kao and Puck, 1968), about 10 years after its first isolation from the Chinese hamster ovaries. Later, the DXB11 was generated by mutagenesis targeting the dihydrofolate reductase gene (*Dhfr*) from K1 cells, resulting in missense mutation in one allele and complete deletion in the other (Urlaub and Chasin, 1980; **Figure 1a**). Note that the *Dhfr* gene is responsible for reducing folate, which is an essential step in nucleotide biosynthesis. Even though both alleles of the *Dhfr* gene were modified in the -DXB11 cells, it still presented some minor activity of the gene. Thus, the deletion of both alleles of this gene led to the generation of the -DG44 cell line in the mid-1980s (Urlaub et al., 1983). Here, it should be noticed that DG44 was not derived from the K1 cell line, but from the very first isolated CHO cell lines (**Figure 1a**). Interestingly, although these cell lines are derived from a common ancestor, extensive mutagenesis and the clonal diversification has resulted in considerable genetic heterogeneity as revealed by their genome sequences (Kaas et al., 2015; Lewis et al., 2013; Xu et al., 2011). Significant differences in the gene copy numbers, single nucleotide polymorphisms, structural variants and chromosomal rearrangements were observed.

**Figure 1.**
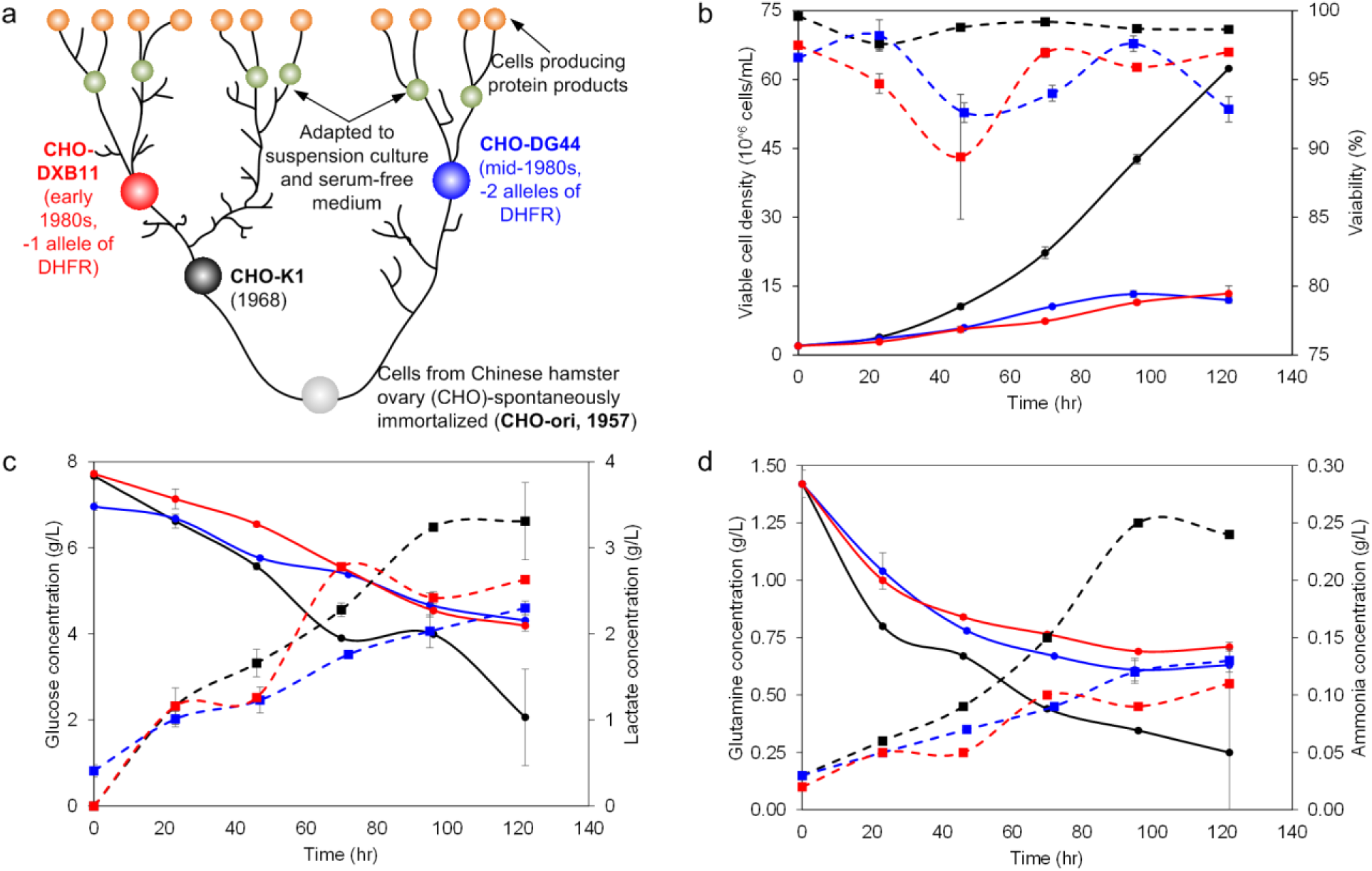
Genetic and phenotypic differences of CHO parental cell lines. a) The known genetic background of the three CHO cell lines studied, i.e. CHO-K1, DG44 and DXB11 (figure re-drawn based on Wurm and Hacker, 2011), b) viable cell densities and % viabilities of three parental cell lines grown in suspension cultures, c) residual concentrations of glucose and lactate, and d) concentration of glutamine and ammonia in the three cell cultures. The black, blue and red lines in each graph represent the CHO-K1, DG44 and DXB11 cell lines, respectively.

Due to the presence of such “quasispecies” within the CHO parental cell lines (Wurm, 2013), there exist considerable dissimilarities in their phenotypes and possibly even variations in the critical quality attributes, e.g. N- and O-glycosylation profiles, of the recombinant proteins produced from them. For example, when the same antibody was produced from different CHO cells, the viable cell densities and final titer across several bioprocess conditions varied significantly, depending on the parental cell line used (Hu et al., 2013; Reinhart et al., 2018). Furthermore, there are appreciable variations among the N-glycosylation patterns and other quality attributes of the recombinant proteins produced (Könitzer et al., 2015; Reinhart et al., 2018). Therefore, it is highly required to understand the diversity in the phenotypes of CHO cells and associate it with known genetic heterogeneity for the rational selection of suitable parental cells. Moreover, the characterization of N- and O-glycosylation preferences of various CHO host cells is a critical aspect within the “Quality by Design (QbD)” framework (Butler and Spearman, 2014).

Towards the goal of unraveling the heterogeneity of the various CHO parental cell lines and linking it with their genetic diversity, here we have accomplished an integrative multi-omics analysis, at the transcriptomic, proteomic and glycomic levels, of the K1, DG44 and DXB11 cells during the exponential growth phase in suspension cultures. To our knowledge, although a recent study reported proteomics analysis to compare the differential protein expression in CHO-K1, -S and *Dhfr*^−^ cell lines (Xu et al., 2017), the current work provides more comprehensive multi-omics analysis for various CHO host cells, thus serving as a basis to further rationalize cell line development and engineering.

## 2 MATERIALS AND METHODS

### 2.1 Cell lines, cultivation and characterization

CHO-K1 (ATCC No. CCL-61, American Type Culture Collection, USA), CHO-DXB11 (ATCC No. CRL-9096, American Type Culture Collection) and CHO-DG44 (Gibco™ catalog number 12609-012, Invitrogen, USA) previously adapted to serum free suspension culture were propagated in a DMEM/F12-based protein free chemically defined medium (PFCDM) supplemented with Soybean peptone (Catalog number P0521, Sigma-Aldrich, USA) and HT supplement (Gibco™ catalog number 11067-030, ThermoFisher Scientific, USA). To compare cell growth over 6 days, all three cell lines were cultivated in batch mode by seeding 2×10^5^ cells/mL into orbitally agitated disposable Erlenmeyer flasks (Corning, USA) in duplicates. Replicate shake flasks were cultivated similarly and harvested at Day 2 for the various –omics analyses. Cell density and viability were measured daily by the trypan blue dye exclusion method using Vi-Cell XR (Beckman Coulter, USA). Glucose, lactate, ammonium, glutamine and glutamate concentrations in the culture supernatant were measured daily using BioProfile 100 Plus (Nova Biomedical, USA).

### 2.2 Analysis of extracellular nutrients/metabolites

Glucose, lactate, glutamine and glutamate concentrations in the culture supernatant were measured using a 2700 Biochemistry Analyser (Yellow Springs Instruments, USA). Ammonia concentration was measured using a commercial kit (Sigma, USA).

### 2.3 Transcriptome profiling and data processing

Cell pellets (1×10^7^ cells) were collected from three replicate flasks for each of the cell lines. Total RNA was isolated using RNeasy Mini Kit (Qiagen, Germany). RNA quality and integrity were verified using a NanoDrop 2000 spectrophotometer (Thermo Fisher Scientific) and Agilent Bioanalyzer 2100 (Agilent Technologies, USA). cDNA targets were generated from 250 ng total RNA using the GeneChip 3’ IVT PLUS Reagent Kit (Thermo Fisher Scientific). The targets were then hybridized onto a proprietary CHO 3’ IVT microarray (CHO V3) consisting of 43,856 probe sets that correspond to 13,358 mouse genes based on CHO EST Sequence and Annotation Release 06 (Affymetrix, Singapore). Quality assessment and data analysis were carried out on the CEL files obtained, using Partek Genomics Suite version 6.6 (Partek Inc., USA).

### 2.4 Proteome profiling and data processing

1×10^7^ cells were collected for each experimental condition in set of biological triplicates. Cells were washed twice in 1 × PBS and lysed in 8 M urea supplemented with 1 × Halt Protease Inhibitor Cocktail (Thermo Fisher Scientific, USA). Insoluble cellular material was removed by centrifugation at 18,000 ×g for 20 min at 4°C, and protein concentration in each sample was determined by Bradford assay using Pierce™ Coomassie Plus Assay Reagent (Thermo Fisher Scientific). Subsequently, 100 μg of protein from each sample was digested with trypsin and labelled with TMTsixplex™ Isobaric Mass Tagging Kit (Thermo Fisher Scientific) according to manufacturer’s instructions. Labelled peptides from each set of replicates were then combined and cleaned up using SCX cartridge system (Applied Biosystems, USA) prior to 2D-LC-MS analysis.

First dimension LC separation consisted of high pH reverse-phase fractionation at a flow rate of 1 ml/min from a gradient of 3.5-54% acetonitrile in 20 mM ammonium formate, pH 9.6 over 30 min on an XBridge BEH C18 column, 130Å, 3.5 μm, 3 mm × 150 mm (Waters, USA) in an Ultimate 3000 UHPLC system (Thermo Fisher Scientific), with a total of 10 pooled peptide fractions from each triplicate set collected. Second dimension separation was carried out by analyzing 3 μg of each sample by nanoLC-MS/MS at a flow rate 300 nl/min 2-50% acetonitrile in 0.1 % formic acid over 100 min on an ACQUITY UPLC Peptide BEH C18 column, 130Å, 1.7 μm, 75 μm × 200 mm (Waters) in a nanoACQUITY UPLC system (Waters, USA) coupled to an LTQ-Orbitrap Velos mass spectrometer (Thermo Fisher Scientific). The LTQ-Pribtirap Velos was operated in data-dependent mode with full mass scan acquired for mass range of 400-1,400 m/z. The top 10 most intense peaks with charged state ≥ +2 were fragmented by HCD with normalized collision energy of 45%, minimum signal threshold of 500, isolation width of 2 Da and activation time of 0.1 ms.

Raw MS data was analyzed by Sequest HT and MS Amanda search engines against NCBI CHO refseq database using Proteome Discoverer v. 2.0 (Thermo Fisher Scientific). Precursor mass range of 350-5,000 Da was used, with MS1 tolerance of 10 ppm and MS2 tolerance of 0.02 Da, static modifications of carbamidomethylation on cysteine and TMTsixplex label on lysine, dynamic modifications of oxidation on methionine, TMTsixplex on peptide N-terminus and acetylation on protein N-terminus, and PSM validation by percolator node. Quantification was performed without replacement of missing value and without application of isotope correction factors. Co-isolation threshold was set at 50% with peptide abundance for each channel normalized by total peptide amount and scaled on channels average.

### 2.5 Metabolome profiling and data processing

1×10^7^ cells were collected from each experimental condition and quenched in 5 volumes of ice-cold 150 mM sodium chloride (Sigma-Aldrich) solution. Subsequently, the quenched cell pellet was processed using a two-phase liquid extraction protocol involving methanol, tricine solution and chloroform as previously described (Yusufi et al., 2017). Polar metabolite extracts were stored at −80°C. Prior to analysis, the extracts were dried and reconstituted in cold methanol and water (5:95 v/v).

Untargeted LC-MS analysis of the polar metabolites was then performed using an ultra-high performance liquid chromatography (UPLC) system (Acquity; Waters, USA) coupled to a mass spectrometer (QExactive MS; Thermo Scientific, USA). A reversed phase (C18) UPLC column with polar end-capping (Acquity UPLC HSS T3 column, 2.1mm, 100 mm, 1.8 mm; Waters) was used with two solvents: ‘A’ being water with 0.1% formic acid (Fluka, Sigma-Aldrich, USA), and ‘B’ being methanol (Optima grade, Thermo Fisher Scientific, USA) with 0.1% formic acid. The LC program was as follows: the column was first equilibrated for 0.5 min at 0.1% B. The gradient was then increased from 0.1% B to 50% B over 8 min before being held at 98% B for 3 min. The column was washed for a further 3 min with 98% acetonitrile (Optima grade, Fisher Scientific) with 0.1% formic acid and finally equilibrated with 0.1% B for 1.5 min. The solvent flow rate was set at 400 mL/min; a column temperature of 30°C was used. The eluent from the UPLC system was directed into the MS. High-resolution mass spectrometry was then performed in both positive and negative electrospray ionization (ESI) modes, with a mass range of 70 to 1,050 m/z and a resolution of 70,000. Sheath and auxiliary gas flow was set at 30.0 and 20.0 (arbitrary units) respectively, with a capillary temperature of 400°C. The spray voltages were 1.25 kV for positive and 1.5 kV negative mode ionization. Mass calibration was performed using standard calibration solution (Thermo Scientific) prior to injection of the samples. A quality control (QC) sample comprising of equal aliquots of each sample was run at regular intervals during the batch LC-MS runs.

The raw LC-MS data obtained was pre-processed based on the XCMS peak finding algorithm (Smith et al., 2006) and the QC samples were used to adjust for instrumental drift. Total area normalization was then applied to the pre-processed data prior to statistical analysis using multivariate (SIMCA-P+ software, version 13.0.3, Umetrics, Sweden) and univariate tools, including relative ratios, student’s t-test (Welch’s correction) and hierarchical clustering for the classification of common trends. Metabolite identities were confirmed by MS-MS spectral comparison with commercially available metabolite standards or with available mass spectral libraries (Wishart et al., 2018).

### 2.6 N- and O-glycome profiling

Glycomics analysis of CHO cells was performed according to a previous study by North et al., (2010) with slight modifications. Briefly, N-glycans were released by PNGase F (Prozyme, USA) from post nuclear fraction of CHO cell lysate which had been subjected to reduction, S-carboxylmethylation, and tryptic digestion. O-glycans were released by sodium borohydride-mediated beta-elimination, from de-N-glycosylated peptides. Both N- and O-glycan samples were permethylated and purified by C18 solid phase extraction. N-glycan samples were analyzed by 5800 TOF/TOF mass spectrometer (AB SCIEX, USA) and O-glycans were analyzed by 5500 Q-TOF system online coupled downstream to a nano-LC system with an integrated chip-based nano C18 column (both from Agilent Technologies, USA). Deconvoluted mass spectra were analyzed for data obtained from both MALDI and QTOF systems. Neutral masses were extracted and parsed against theoretical N- and O-glycan mass libraries (permethylated for N-glycans, reduced and permethylated for O-glycans) for CHO cells. Putative N- and O-glycan structures were assigned based on the knowledge of glycan biosynthetic pathway in CHO cells (North et al., 2010).

### 2.7 Identification of differentially expressed genes in the transcripts and proteins

Using the normalized gene expression data, the differentially expressed genes was identified with CHO-K1 as the reference using empirical Bayes-moderated t-statistics via “limma” package in R (Wettenhall and Smyth, 2004). Genes showing ≥ ±1.5 fold difference with a FDR-adjusted p-values ≤ 0.01 were classified as differentially expressed.

Quantifiable proteins were identified at 1% FDR with ≥ 2 unique peptides and had abundance values reported for at least two of the triplicate samples; any missing protein abundance value was replaced by the average abundance values reported for the other two of the triplicate samples where applicable. Proteins showing ≥ ±1.5 fold difference with a FDR-adjusted p-values ≤ 0.1 were classified as differentially expressed.

### 2.8 Metabolic pathway enrichment analysis of differentially expressed transcripts and proteins

Enrichment analysis of the metabolic genes was performed using omics data in conjunction with the CHO genome-scale metabolic network (Hefzi et al., 2016). The gene-protein-reaction (GPR) associations in the metabolic network and the reaction pathway classifications were used to assign the metabolic genes to different reaction pathways. To identify enriched metabolic pathways, the differentially expressed genes are first identified based on the p-value (<0.01; FDR-corrected) and fold change (>1.5). The enrichment of differentially regulated genes in each pathway was then computed using the hypergeometric distribution as follows:

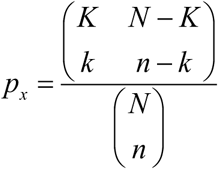

where *N* is the total number of genes in CHO cells, *K* is the total number of up/downregulated genes, *n* is the total number of genes in pathway “*x*” and k is the number of up/downregulated genes in pathway “*x*”. The pathway with lowest *p*-value is classified as highly enriched in differentially expressed genes.

### 2.9 Estimation of gene copy number variations (CNV) between CHO cell lines

The changes in absolute gene copy numbers between the CHO cell lines were calculated using CHO-K1 as the reference from the normalized sequencing read depth of individual cell lines which was reported previously (Kaas et al., 2015). A gene is considered to be deleted in a cell line if the read depth is zero in that particular cell line but >0.95 in CHO-K1. Gene amplifications and reductions were also calculated in a similar manner.

## 3 RESULTS

### 3.1 CHO-K1 grows faster in suspension batch cultures than DG44 and DXB11

The three CHO parental cell lines, i.e. K1, DG44 and DXB11, were propagated in duplicates in the same DMEM/F12-based protein free chemically defined medium (PFCDM) supplemented with Soybean peptone and HT supplement, in order to reduce variability due to differences in cell culture media. These cell culture experiments clearly showed that the K1 cells grew much faster when compared to the other two cell lines, reaching the maximum cell densities first (**Figure 1b**). The doubling time of K1 cells is 20±1 hr whereas DG44 and DXB11 correspond to 30±0.3 and 32±4, respectively. Notably, these results were highly consistent with an earlier study which reported that the maximum viable cell densities achieved in batch culture following the order of K1>>DG44>DXB11 (Sommeregger et al., 2013). While the cells were grown in commercial media in the earlier study, the concordance between the two results highlight that the slow growth of *Dhfr*^-^ cells is mainly because of the genetic divergence. Here, it should be highlighted that although all the three media contained HT, the purine supplements enabling the cells to overcome deprivation of nucleoside precursors as a result of the *Dhfr* deficiency, we still observed much lower growth rates than K1. This confirms that the loss of *Dhfr* gene activity could not be simply compensated by supplementing HT as there could be additional pathways contributing to such growth differences (Florin et al., 2011). In order to further investigate metabolic traits of CHO host cells, we also profiled the major nutrients and byproducts in the cell culture. Glucose was rapidly consumed in the K1 compared to others (**Figure 1c**). Similarly, a sharper decrease in the glutamine concentrations was also observed in K1 cell cultures with a concomitant higher accumulation of ammonia and lactate than DXB11 and DG44 cell cultures (**Figure 1d**), possibly indicating that K1 cells undergo higher carbon and nitrogen catabolism.

### 3.2 Global transcriptome and proteome profiling of CHO parental cell lines confirms Known genetic heterogeneity

To systematically analyze the underlying molecular mechanisms leading to the observed growth phenotypes across CHO host cells, samples were harvested during the exponential phase for transcriptome and proteome profiling. Transcriptomics analysis was performed using a proprietary microarray encompassing CHO probe sets that correspond to 13,358 unique mouse genes (**Supplementary file**). As expected, the expression of *Dhfr* gene in K1 was more than twice the intensity of DXB11 while a very low detectable signal was present in DG44, possibly due to noise, thus confirming the known genetic backgrounds of CHO cells. MS-based proteomic profiling using multiplexed isobaric tags (iTRAQ) identified and detected a total of 1998 unique proteins corresponding to 1983 genes in more than one biological replicate (**Supplementary file**). Note that 1857 of these 1983 genes were also present in the microarray, thus providing relevant information at both the transcript and protein levels (**Figure 2a**). After transcriptome and proteome profiling, we initially analyzed the global mRNA and protein expression patterns via two different methods: 1) principal component analysis (PCA) and 2) hierarchical clustering of Pearson correlation coefficients. PCA revealed that the expression patterns of all the cell lines were markedly different at both transcript and protein levels (**Figure 2b and 2c**), highlighting that the genetic divergence of the CHO parental cell lines indeed influences the phenotype via transcriptional and translational regulations. The hierarchical clustering of transcriptome and proteome also showed similar trends, where the global expression patterns of K1 are closer to DXB11 than DG44 cells at both transcript and protein levels (**Figure 2d and 2e**), recapitulating the known genetic divergence of the CHO parental cell lines (**Figure 1a**).

**Figure 2.**
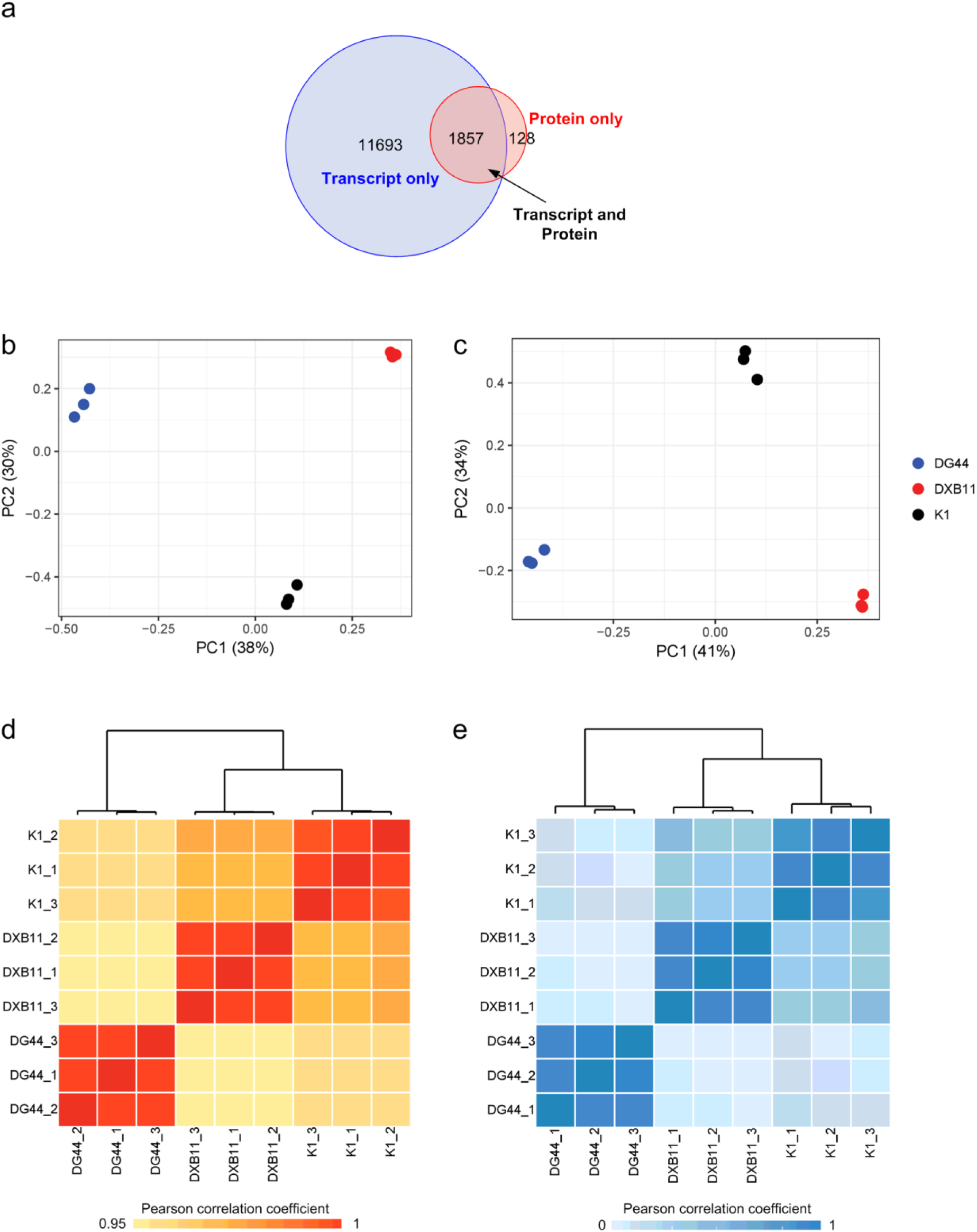
Global mRNA and protein expression patterns of CHO parental cell lines. a) Global coverage of transcriptome and proteome data, b) PCA of the transcriptome, c) PCA of the proteome, d) hierarchical clustering of transcriptome and e) hierarchical clustering of proteome. Clustering was based on Euclidean distance of Pearson correlation coefficients.

### 3.3 Integrative transcriptome and proteome analysis highlights that cellular metabolism and cell cycle progression are negatively regulated in *Dhfr*-deficient cells

To pinpoint relevant cellular functions attaining the observed phenotypic differences based on the transcriptomic and proteomic features, we first identified the differentially expressed genes at both levels using K1 as a reference. In the K1 vs DXB11 comparisons, we identified 2231 mRNAs and 239 proteins to be differentially expressed, with a large fraction being down-regulated in DXB11 (**Supplementary file**). A total of 2875 mRNAs and 304 proteins were also differentially expressed in K1 vs DG44 comparisons where some are down-regulated in DG44. Together, the larger fraction of up-regulated compared to down-regulated genes in K1 suggests more active transcription and translation compared to DG44 and DXB11. Subsequent gene ontology-based enrichment analysis of the differentially expressed transcripts and proteins revealed that the cellular metabolic process including lipid and energy metabolism, DNA, RNA and protein metabolic processes, cell cycle, growth, and cell proliferation and death present the major expression differences among the three cell lines (**Figure 3a**). Particularly, both DG44 and DXB11 presented a negative regulation of cell cycle progression, DNA replication and the biosynthesis of nucleotides. Several genes which commit the cells in cell cycle from G2 phase to M phase and the next mitotic phase were all down-regulated in DG44 and DXB11. Further, the down-regulation of cyclins including *Ccny*, *Ccnl2*, *Ccnc* and *Ccnh*, and serine/Threonine Kinase, *Akt1*, indicates that the cell cycle progression and DNA replication are negatively regulated by PI3K-Akt and MAPK signaling pathways (Duronio and Xiong, 2013; Rhind and Russell, 2012). In addition to the cell cycle and cellular organization related genes, we also found that the purine nucleotide metabolic processes (as expected, due to the *Dhfr* gene copy number differences) and sterol metabolic process to be significantly enriched in the differentially expressed genes. Remarkably, the expression patterns of glycosylated protein processing processes such as N-glycosylation, Golgi organization, vesicle-mediated protein transport and protein folding were also significantly altered across cell lines.

**Figure 3.**
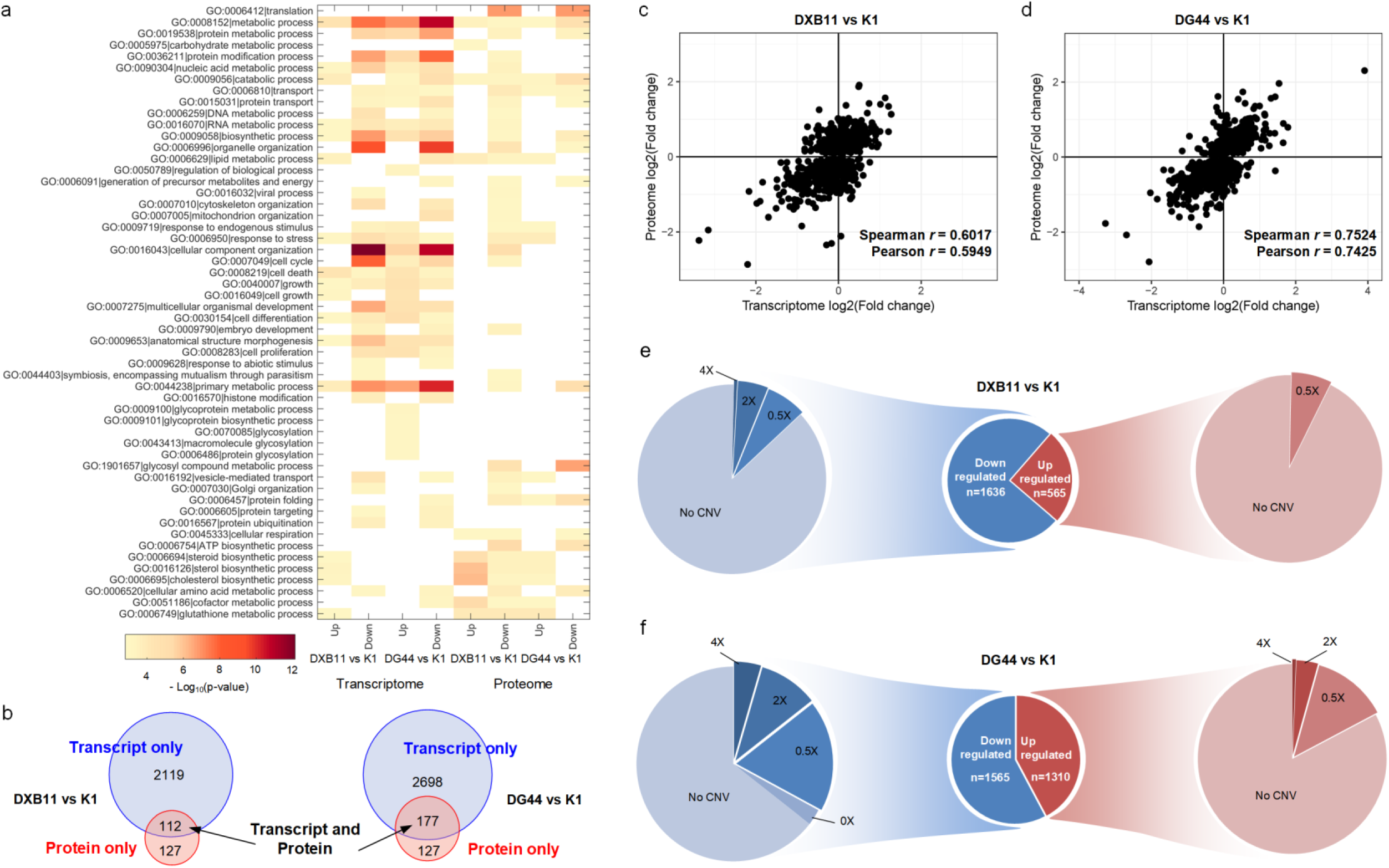
Differential expression signatures of CHO parental cell lines. a) GO enrichment analysis of differentially expressed genes at both transcript and protein level, b) number of differentially expressed genes at transcript and protein level in DXB11 and DG44 w.r.t. K1, c) correlation between transcript and protein expression ratios in DXB11 vs K1, d) correlation between transcript and protein expression ratios in DG44 vs K1, e) the effect of genomic variations in differentially expressed genes at the transcript level in DXB11 vs K1 comparison and f) the effect of genomic variations in differentially expressed genes at the transcript level in DG44 vs K1 comparison.

We next overlaid the transcriptomic and proteomic expression patterns to compare their concordance in each comparison (**Figure 3b**). A set of 55 proteins commonly downregulated in DG44 and DXB11 at both levels were identified, including *Pacsin2* and *Ezr*, both of which are substrates of protein kinases associated with the regulation of actin cytoskeleton dynamics (D’Angelo et al., 2007; Goh et al., 2012; de Kreuk et al., 2011), altering the cellular structure as well as cell migration and macromolecular assembly. The current analysis also identified 110 and 113 up- and down-regulated proteins in DXB11 vs K1 and DG44 vs K1 comparisons, respectively, which do not show any expression differences in their corresponding mRNAs (Supplementary file). Their gene enrichment analysis allowed us to understand that central metabolic pathways, particularly energy metabolism including the ATP biosynthesis (e.g., *Atp5j*, *Atp5o* and *Atpif1*) via cellular respiration and ROS detoxifying glutathione metabolism, are post-transcriptionally regulated. In addition, some of the genes from nucleotide metabolism and glycosyl compound biosynthetic pathways were also potentially post-transcriptionally regulated. Such examples include *Ada*, *Neu2*, *Galnt2* and *Rrm2*.

The fold-changes of 1857 proteins that are detected at both protein and transcript levels showed a moderate correlation with corresponding transcript fold-changes in both DXB11 vs K1 (Spearman’s *r* = 0.6017; Pearson’s *r* = 0.5949; **Figure 3c**) and DG44 vs K1 (Spearman’s *r* = 0.7524; Pearson’s *r* = 0.7425; **Figure 3d**). Previously, several studies have reported similar correlations (moderate or poor) which are attributable to possibly the changes in translation and transcription rates, protein and transcript degradation rates and other regulatory factors (Liu et al., 2016). However, despite the moderate correlation, we were able to identify a list of 109 genes from such comparisons as potential targets for cellular engineering in CHO cells since they showed a very good correlation in expression between transcript and protein levels (R^2^ > 0.9; **Supplementary file**).

### 3.4 Hard-wired changes of gene expression in signaling and nucleotide metabolic pathways are observed in DXB11 and DG44 cell lines

In order to understand whether significant differences in the transcriptional and translational patterns across the cell lines are caused by the genetic heterogeneity, we examined their gene copy number changes. Interestingly, this analysis showed that 9% of the total 2231 differentially expressed mRNAs have a hard wiring of copy number changes in the DXB11 genome when compared to K1 (**Supplementary file**; **Figure 3e**). 51 up-regulated genes have increased copy numbers including two serine/threonine-protein kinases (*Wnk2* and *Mknk2*) that are known to inhibit cell proliferation by negatively regulating the ERK pathway (Moniz et al., 2007); 154 down-regulated genes showed a decrease in gene copy numbers, including homozygous deletion of two genes, *Tmem14a* and *Plgrkt*. Notably, *Tmem14a* is a transmembrane protein which inhibits apoptosis by negatively regulating the mitochondrial membrane permeabilization (Woo et al., 2011). Therefore, its complete deletion in DXB11 could possibly result in dysregulation of apoptosis inhibition pathway, contributing to the cell death. In addition, certain cell cycle associated genes such as *Cdk11b*, *Taf4* and *Dscc1* were also heterozygously deleted, presumably leading to the negative regulation of cell cycle progression as mentioned earlier.

When compared to DXB11 cells, even larger fraction of differentially expressed mRNAs (18%) are associated with the copy number changes in the DG44 genome (**Figure 3f**). Interestingly, DG44 cells showed homozygous deletion of 12 genes including the known removal of *Dhfr*. In addition, passenger deletion of neighboring genes (*Fam15b*, *Msh3* and *Zfyve16*) and other cell cycle related genes (e.g., *Cdk11b*, *Taf4* and *Dscc1*) were observed. Clearly, these results suggest that DG44 cell lines have the inherent limitation in cell cycle progression and nuclear division as reflected in the slow growth phenotype. Several up-regulated genes in DG44 also have corresponding augmentation in gene copy numbers where some genes were amplified up to 4 times. Such examples include vesicle-mediated transport regulating genes, *Axl* and *Synj2bp*, which are related to the vascular transport of proteins. The complete list of differentially expressed genes with corresponding copy number differences are provided in Supplementary file.

### 3.5 Dysregulation of nucleotide and sterol metabolism may lead to slow growth in *Dhfr*-deficient cell lines

Since majority of the differentially expressed genes were from cellular metabolism, we integrated the transcriptome and proteome data with the CHO genome-scale metabolic network (Hefzi et al., 2016) to unravel the transcriptional signatures of the CHO metabolism. The transcriptome and proteome data were mapped onto the metabolic network using the available gene-protein-reaction (GPR) rules. The microarray-acquired transcriptome data maps to 1287 of 1764 gene loci in the model while the MS-acquired proteome data is linked to 548 genes. Following the transcriptome and proteome data mapping, we assessed the enrichment of various metabolic pathways among the up-/down-regulated transcripts and proteins (see **Methods**; **Figure 4a**). As expected, the purine nucleotide metabolism is grossly dysregulated in both DG44 and DXB11 when compared to K1 due to the absence of *Dhfr* gene. Particularly, adenylosuccinate lyase (*Adsl*), an enzyme which converts the succinylaminoimidazole carboxamide ribotide (SAICAR) into key AMP precursor, aminoimidazole carboxamide ribotide (AICAR), was downregulated by more than 3.5 times in both cell lines (**Figure 4b**). Similarly, nicotinamide nucleotide adenylyltransferase 2 (*Nmnat2*) which catalyzes an essential step in NAD biosynthetic pathway was also downregulated by more than 3 times in both cell lines. Apart from nucleotide metabolism, interestingly, several genes in the cholesterol biosynthesis including *Hmgcs1*, *Fdps*, *Idi2*, *Lss*, *Ebp* and *Hsd17b7* were up-regulated in DG44 and DXB11 (**Figure 4b**). Since endogenous cholesterol levels modulate the epidermal growth factor receptor (EGFR) signaling and control meiosis (Gabitova et al., 2014), elevated sterol levels in *Dhfr*-deficient cell lines could have a potential role in negatively regulating cell cycle progression although it is unclear how exactly sterols control the cell cycle. ROS-detoxifying glutathione metabolism was observed to be upregulated, possibly suggesting that these cell lines have better ability for protein processing as an earlier study reported that higher levels of glutathione were found in high-producer when compared to low-producer CHO cells (Orellana et al., 2015).

**Figure 4.**
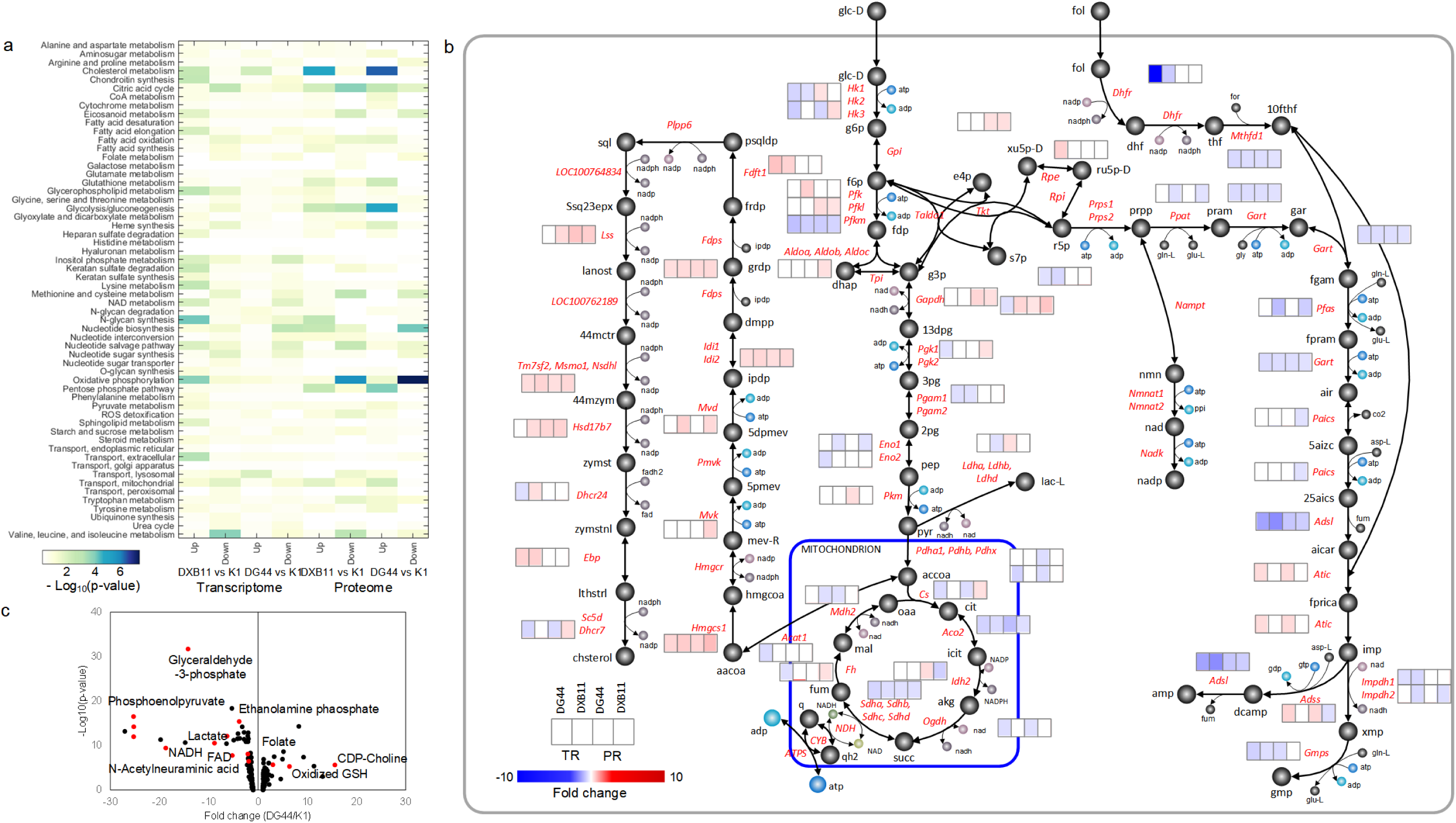
Differential expression signatures of metabolic genes in CHO parental cell lines. a) The enrichment of various metabolic pathways in the CHO genome-scale network at both transcript and protein level and b) the differential expression of various genes from central metabolism, de novo purine biosynthesis and sterol metabolism at transcript and protein levels are shown in the metabolic map, and c) comparison of intracellular metabolite levels in DG44 and K1.

To further examine whether such observed expression differences in the transcripts and proteins of metabolic pathways ultimately affect the endogenous metabolite levels, we performed LC-MS based metabolite profiling of DG44 and K1 cells. Noticeably, this analysis showed several metabolites in central metabolism, e.g., glyceraldehyde-3-phosphate and lactate, and energy generation (NAD and FAD), are much abundant in K1 (**Figure 4c**). On the other hand, the levels of metabolites that are associated with cellular stress and growth inhibition such as oxidized glutathione and CDP-choline were observed to be higher in DG44 (**Supplementary file**).

### 3.6 N- and O-glycosylation of host cell proteins from parental CHO cell lines is discernably different

The considerable genetic heterogeneity of various parental CHO cells is also thought to influence the quality attributes such as N- and O-glycosylation, charge variants and sequence variants of the recombinant proteins produced from them. Therefore, in order to analyze the possible variations in quality attributes of the protein products, we profiled the N- and O-glycosylation of the host cell proteins. Corresponding glycomic analyses successfully detected 38 different N-glycan species which are usually processed in the ER, cis-Golgi, medial-Golgi and trans-Golgi compartments (See Methods). Comprehensive analyses of the resulting N-glycan species and their relative abundances revealed several interesting aspects. The initial processing in ER is not likely to be different among the cell lines; the high mannose structures were found to be relatively similar. However, there exists an appreciable difference in downstream processing of N-glycans. The terminal GlcNAc containing hybrid structures processed at the cis-Golgi were slightly higher in DXB11 than other two cell lines while the complex tri-antennary structures which are processed in medial- and trans-Golgi were higher in DG44 and K1 (**Figure 5a**). Notably, K1 cells had much higher final capping of glycan species, i.e. sialylation, beyond which no further sugar moieties can be added. Apart from N-glycans, we also measured the relative abundances of O-glycans including three major structures, T antigen, sialyl-T antigen and diasialyl-T antigen. Further, the sialylated structures were found to be more abundant in K1 than the other two cell lines even in O-glycans (**Figure 5b**).

**Figure 5.**
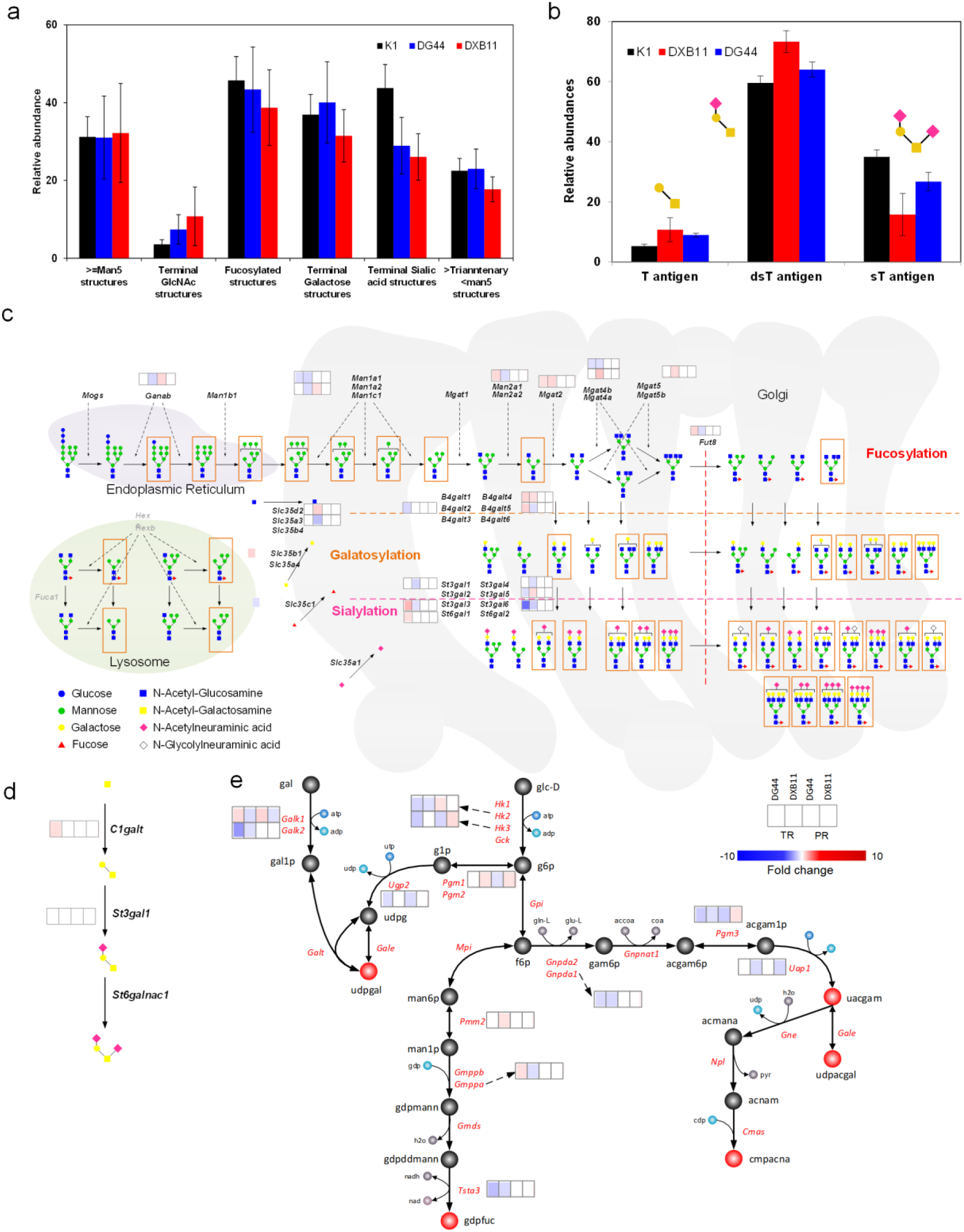
Glycosylation differences between the CHO parental cell lines. a) N-glycosylation differences, b) O-glycosylation differences, c) the differentially expressed genes and corresponding glycan species in N-glycosylation pathway, d) differentially expressed genes in O-glycan pathway and e) the biosynthetic pathway of nucleotide-sugar donors. The glycans enclosed in boxes in (c) were detected in our experiments.

We subsequently correlated the variations in N-glycan species with the transcriptomic and proteomic data of the N-glycosylation associated pathways (**Figure 5c**). Although there was no appreciable difference in the expression of genes in early processing of N-glycans, one notable exception is that the family of N-acetylglucosaminyltransferase genes (*Mgat2*, *Mgat4b* and *Mgat5*) were up-regulated in DXB11. Consistent with this observation, the terminal GlcNAc structures were slightly abundant in DXB11. In addition, the high levels of terminal galactose structures were correlated well with the up-regulation of two beta-1,4-galactosyltransferase isoforms (*B4galt4* and *B4galt5*), the enzymes which add galactose to the GlcNAc structures, and the UDP-galactose transporter (*Slc35b1*). Similarly, four isoforms of beta-galactoside alpha-2,3-sialyltransferase (*St3gal1*, *St3gal2*, *St3gal5* and *St3gal6*) were all up-regulated in K1, supporting the high levels of CMP-sialic acid structures (**Figure 4c**).

## 4 DISCUSSION

Currently, 7 out of the top 10 drugs are industrially produced from CHO cells where various companies use different CHO host cell lines empirically based on their experience and regulatory track records. While it has been argued that the possible differences in the CHO host cells could present appreciable variations in therapeutic protein production which could be further complicated by the clonality, still the literature available on comparing various CHO hosts is relatively scarce. Therefore, in order to evaluate the phenotypic traits and bioprocessing potentials of CHO host cell lines and provide a basis for rational cell line selection, we compared the three most widely used CHO cell lines via multi-omics profiling for the first time ever in this study. As such, our analysis uncovered that the *Dhfr*^-^ cell lines, i.e. DG44 and DXB11, grow much slowly due to the mutations in purine nucleotide biosynthesis pathways, which retarded the cell growth directly as well as indirectly by interfering with DNA replication and cell cycle progression. Interestingly, this study also showed that the passenger deletions in both *Dhfr*^-^ cell lines have critical impact on cell cycle progression and growth factor signaling as the loss or gain of gene functions associated with these pathways reflected well at the transcriptomic and proteomic levels. Moreover, the N- and O-glycomic profiling of host cell proteins in all the three cell lines revealed notable differences in the resulting glycosylation, thus highlighting the need to select a relevant cell line for the target protein of interest based on the preferred glycosylation patterns.

Consistent to earlier reports (Florin et al., 2011; Sommeregger et al., 2013), our study confirmed the ability of K1 cells to grow faster than the DG44 and DXB11 under suspension cultures. The transcriptomic and proteomic profiling uncovered that this physiological difference could be a consequence of two scenarios: 1) the lack of *Dhfr* gene leading to severe limitation for nucleotide synthesis and/or 2) the negative regulation of cell cycle via multiple signaling pathways including PI3K-Akt, MAPK and ERK pathways, and a possible pathway activated by endogenous sterol levels (Gabitova et al., 2014). In this sense, overexpression of the several down-regulated cyclins such as *Ccny, Ccnl2, Ccnc and Ccnh*, and serine/threonine kinase, *Akt*, could be an interesting strategy for improving the cellular growth in *Dhfr*^-^ cell lines as it has been shown earlier in several mammalian cells (Jaluria et al., 2007).

In addition to the inhibition of cell cycle progression, several genes associated with cellular macromolecular assembly and component biogenesis were down-regulated in both DG44 and DXB11. Here, it should be highlighted that such observations of varied cellular structure is in good agreement with earlier studies which reported the size of ER and mitochondrial mass in K1 is higher than that of DXB11 (Hu et al., 2013). From a bioprocessing point of view, a larger ER provides significant advantage for K1 cells as it could handle much more proteins for their proper folding and further processing than the other two cell lines. The limitations in the ER size could also be a possible reason for growth inhibition in *Dhfr*^-^ cells; since the ER size is limited, as the unprocessed proteins accumulate, the unfolded protein response pathways could be activated, which in turn could trigger the ROS stress pathways, thus ultimately invoking the apoptosis pathway (Hu et al., 2013). The multi-omics data generated in this study supports this hypothesis at several levels: both transcriptome and proteome showed the up-/down-regulation of several apoptosis activators and inhibitors, respectively. Subsequent metabolome analyses confirmed the high levels of oxidized glutathione, a ROS stress marker, possibly highlighting the relevance of elevated ROS stress in DG44 than K1. However, whether all such pathways operate in tandem has to be confirmed with further experiments.

Our study pinpointed that the N- and O-glycosylation patterns of protein products could have cell line specific variation due to the inherent characteristics of CHO hosts. For example, DXB11 have high amounts of less processed structures, particularly terminal GlcNAc glycans, and K1 show much more sialylated structures in both N- and O-glycosylation profiles, which is possibly attributed to high expression of relevant biosynthetic enzymes as well as high levels of precursor availability. Notably, these observations are consistent with an earlier study which compared the glycosylation profile of an antibody produced from DG44 and K1 cells and observed that K1 produced much higher sialylated structures (Reinhart et al., 2018). While we observed no gene copy number difference in the glycosylation biosynthetic pathway, all the cell lines have their own expression preferences in each step of the glycan processing as different isoforms were up-/down-regulated. Such variation in isoforms expression could possibly be a result of differences in SNPs (Lewis et al., 2013) or due to changes in epigenetic control which await experimental verification. Although the variations of glycosylation patterns can be modulated to a certain degree by modifying the media and feeding strategies, the selection of an appropriate host depending on the product of interest would nonetheless be relevant to achieve consistent quality. To this end, K1 and DG44 are probably more suitable for producing erythropoietin (EPO) or interferon beta 1 (IFNB1) as these molecules have complex tetra-/tri-antennary N-glycan species whereas DXB11 could provide a minor advantage for producing monoclonal antibodies since they have higher terminal GlcNAc structures that are usually bi-antennary.

In summary, our work exploited the wealth of multi-omics technologies to systematically link the possible phenotypic differences across CHO host cell lines with their genotypes. It also clearly highlighted that the genetic heterogeneity of CHO cells indeed affects their growth phenotype as well as N- and O-glycosylation profiles which could impact the bioprocessing steps. Although the integration of transgene could further alter the genomic landscape of host cells compared here, particularly DG44 and DXB11 due to the additional selection step, we believe that most results unraveled in this study should hold good since the transgene targets only fragile regions of the genome and importantly the glycosylation profiles remain unaltered (Yusufi et al., 2017). Therefore, the current study could well serve as a good starting point for the rational selection of relevant CHO host cells as well as to establish cell line engineering targets.

## Acknowledgements

This work was supported by the Biomedical Research Council of A*STAR (Agency for Science, Technology and Research), Singapore, and the Next-Generation BioGreen 21 Program (SSAC, No. PJ01334605), Rural Development Administration, Republic of Korea.

